# A non-coding SNP in *ELF3* alters expression of *ELF3β* and confers adaptation of Arabidopsis to a continental climate

**DOI:** 10.1101/2025.10.23.684042

**Authors:** Christopher R. Buckley, Alexandre Fournier-Level, Michael J. Haydon

## Abstract

Circadian clocks are biological timekeepers that influence most aspects of plant biology and allow organisms to predict and adapt to daily and seasonal environmental cycles. In particular, the important role of circadian clocks in phenology has led to numerous examples of both functional variation in circadian clock genes within natural and domesticated plant populations. To assess variation of circadian rhythms in natural Arabidopsis populations, we have phenotyped 287 accessions using a seedling transformation protocol with a circadian luciferase reporter. A genome wide association study of circadian period identified multiple single nucleotide polymorphisms (SNPs) in the core clock gene *EARLY FLOWERING 3 (ELF3).* We identified three *ELF3* haplogroups that are associated with seasonal variability in temperature found in continental climates and found strong evidence of a selective sweep coinciding with the most recent de-glaciation period in Europe. One of the SNPs is located within intron 2 of *ELF3* in the region upstream of an alternative transcription start site affecting expression of a shorter *ELF3β* transcript. Our results indicate the important role of subtle variation in a core circadian clock gene in adaptation to changing climate.

## Introduction

Terrestrial organisms are exposed to dramatic daily and seasonal changes in their environment. Plants need to anticipate and respond to these changes in a timely manner. This timely responsiveness is particularly important for annual plant species like *Arabidopsis thaliana* (Arabidopsis), where monocarpic reproduction requires that an individual must survive the climate conditions to transmit genes to the subsequent generation. To respond to predictable environmental changes, plants have evolved circadian clocks that synchronise their internal biology with the external cycle. The circadian clock regulates at least 30% of gene expression (Romanowski et al., 2020) and its influence extends to all aspects of plant biology including development, growth, metabolism and physiology (Sanchez and Kay, 2016). Synchronisation of circadian rhythms with environmental cycles confers a fitness advantage to plants by promoting faster growth, higher survival and are increased fecundity (Green et al., 2002; Michael et al., 2003; Dodd et al., 2005; Oravec and Greenham, 2022).

The circadian oscillator integrates environmental cues to calibrate the internal biological time. The current model of the Arabidopsis oscillator is composed of multiple interlocking loops of transcriptional-translational feedback of ∼20 core transcription factors (TFs) (McClung, 2019; Haydon et al., 2019). This oscillator can be simplified to three, core regulatory modules. A group of Myb-like transcription factors including *CIRCADIAN CLOCK-ASSOCIATED 1* (*CCA1*), *LATE ELONGATED HYPOCOTYL* (*LHY*) and *REVEILLEs* (*RVEs*) are expressed in the morning (Alabadí et al., 2001; Farinas and Mas, 2011). This is followed in the daytime by a wave of PSEUDO-RESPONSE REGULATOR (PRR) activity (Matsushika et al., 2000). Finally, a repressive EVENING COMPLEX (EC) of at least three regulatory proteins is active after dusk (Nusinow et al., 2011). Each tightly controlled component of the oscillator regulates hundreds of targets in downstream pathways, the overall effect of which generates circadian rhythms of cellular and physiological activity (McClung, 2019).

Genetic variation in oscillator genes has been shown to disrupt the tightly interacting network and modify parameters of circadian rhythms (Li et al., 2025). Two circadian rhythm parameters, period and phase, are most commonly affected by genetic variation within the oscillator. Period is the length of a full cycle, and phase is the time of peak activity. For example, plants with loss-of-function mutations in *CCA1* or *TIMING OF CAB EXPRESSION 1* (*TOC1*) oscillate with a shorter period (Millar et al., 1995; Green and Tobin, 1999). Changes in the pace of rhythms are typically accompanied by a shift in phase, but phase changes can also arise independently of period (Eriksson and Millar, 2003). Severe genetic defects, like knockout of individual components of the EC, can result in complete arrhythmia (Hicks et al., 1996; Doyle et al., 2002; Hazen et al., 2005).

Changes in circadian rhythm parameters might enable adaptation to different environmental variables, such as photoperiod, seasonality or climate. The circadian oscillator is entrained by fluctuations of light and temperature that vary along geographical and bioclimatic gradients. Notably, the maximum photoperiod differs by almost 12 h over the natural latitudinal range of Arabidopsis (0° to 68° N; Hannah et al., 2006). Thus, differences in the architecture and function of the circadian oscillator are likely required for adaptation across different environments. Latitudinal clines in circadian period have been identified in several annual plants including Arabidopsis (Michael et al., 2003) soybean (*Glycine max*) and monkeyflower (*Mimulus guttatus*) (Greenham et al., 2017). Similarly, domestication and breeding of tomato (*Solanum lycopersicum*) at higher latitudes has slowed its cycling pace (Müller et al., 2016). In addition to daylength, other selection pressures likely act on the oscillator as plant species expand their ranges. The circadian period of Swedish Arabidopsis accessions is not correlated with latitude. Instead, period is more strongly associated with variation in *COLD REGULATED 28 (COR28)*, a gene implicated in the control of cold tolerance and flowering time (Rees et al., 2021).

In addition to genetic variation within the core oscillator, modulation of input pathways, including light signalling (Oakenfull and Davis, 2017) and metabolism (Buckley et al., 2023) can also alter circadian output. Thus, the genetic basis of variation in circadian rhythms between natural populations could be in components outside of the core oscillator. Although loci that affect circadian rhythm parameters have been uncovered from QTL mapping studies (Darrah et al., 2006; Anwer et al., 2014; Rubin et al., 2019), the narrow diversity of biparental mapping populations precludes species-level generalisation. Genome-wide markers have been developed for over 1000 natural Arabidopsis accessions enabling genome-wide association studies (GWAS) (Alonso-Blanco et al., 2016). A previous study of circadian rhythms of leaf movement across 150 Arabidopsis accessions was performed prior to the development of this resource (Michael et al., 2003) and thus has limited overlap with the 1001 Genomes accessions. A more recent study of circadian rhythms of delayed leaf fluorescence in 191 accessions was restricted to a specific geographical range (Rees et al., 2021).

Here, we aimed to assess a fuller genetic picture of adaptive variation in circadian rhythms between natural populations of Arabidopsis. Circadian rhythms were measured using a high-throughput seedling transformation assay with a luciferase reporter in a geographically broad collection of Arabidopsis accessions. GWAS were used to uncover candidate genetic variants responsible for changes in circadian rhythms. We identified a strong association of circadian period with a region surrounding the core circadian clock EC gene *EARLY FLOWERING 3 (ELF3)* and defined three *ELF3* haplogroups that contribute differentially to circadian period, are associated with adaptation to continental climates and confer responsiveness to temperature. Among these is a non-coding genetic variant within an intron of *ELF3* that affects expression of an alternative transcript. We show that variation in *ELF3* has been selected for and probably aided the spread of Arabidopsis to continental climates. Our results highlight the important role of *ELF3* and the circadian clock in adaptation to climate.

## Results

### Extensive natural variation of circadian rhythms in Arabidopsis accessions

The preferred method for accurately measuring circadian rhythms in Arabidopsis is by transcriptional circadian luciferase reporters. Previous surveys of circadian rhythms in natural populations of Arabidopsis have used leaf movement (Michael et al 2003) or delayed fluorescence (Rees et al. 2021), but these are comparably less precise. We used a seedling transformation protocol (Ting et al., 2022) with a circadian clock reporter *GIGANTEAp:LUC2* (*GIp:LUC2*) (Supplementary Figure S1; Supplementary Table S1) to measure circadian rhythms in 287 natural accessions, representing the full geographic and climatic species range of Arabidopsis (Figure 1A-H) (see Materials & Methods). The circadian rhythm data generated from the accessions were highly robust: mean relative amplitude error (RAE) of all accessions was 0.21±0.13, suggesting high quality rhythms across the data set (Figure 1D; 1H). We detected significant variation for period and phase of luciferase rhythms. The mean period was 21.1±1 h (Figure 1A) ranging from 19.0 h in the Russian accession Lebja-1 and 24.8 h in T570 from Sweden (Figure 1E). Period was strongly genetically determined, with a broad-sense heritability (H^2^) of 0.52. We did not detect a correlation between period and latitude (Supplementary Figure S2A) or between period and longitude (Supplementary Figure S2B) in our data. The mean phase across accessions was 14.4±2 h after subjective dawn (Figure 1B), which is consistent with the expected dusk-phased expression of *GI*. Broad sense heritability was somewhat lower for phase than period (H^2^ = 0.28), but more than 7 h difference was found between the earliest phase of GIp:LUC2 in Tul-0 and the latest phase in Lerik1-3 (Figure 1F). The amplitude of rhythms also showed large variation (Figure 1C,G), although the broad sense heritability of this trait was relatively low (H^2^ = 0.15).

**Figure 1:**
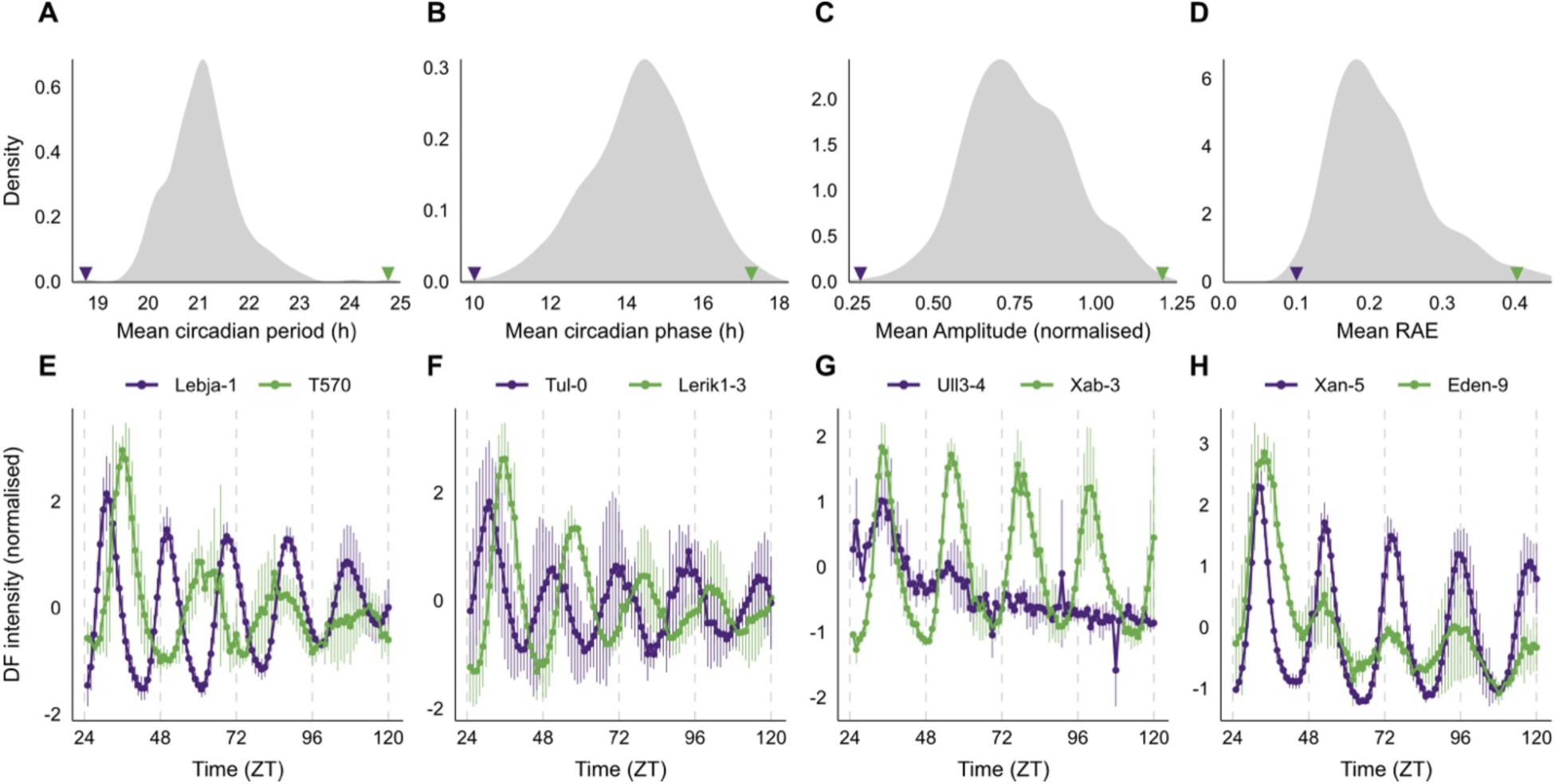
Variation in circadian rhythms of natural Arabidopsis accessions. Circadian rhythms were measured in 287 *Arabidopsis thaliana* accessions using seedling transformation of *GIp:LUC2*. Distributions of **A,** circadian period **B,** circadian phase, **C,** amplitude and **D,** relative amplitude error (RAE). Purple and green arrows indicate position in the distribution of example accessions shown in **E**, **F**, **G** and **H**. Values are mean ±sd, n=4.

### A genomic region containing ELF3 is associated with variation in circadian period

To identify the genetic underpinnings of variation in circadian rhythms, we performed GWAS. No SNPs were detected as significantly associated with circadian phase, amplitude or RAE (Supplementary Figure S3). However, we identified 383 SNPs significantly associated with circadian period (Figure 2A; Supplementary Table S2). These SNPS were located within the coding sequences of 43 genes (Figure 2A).

**Figure 2:**
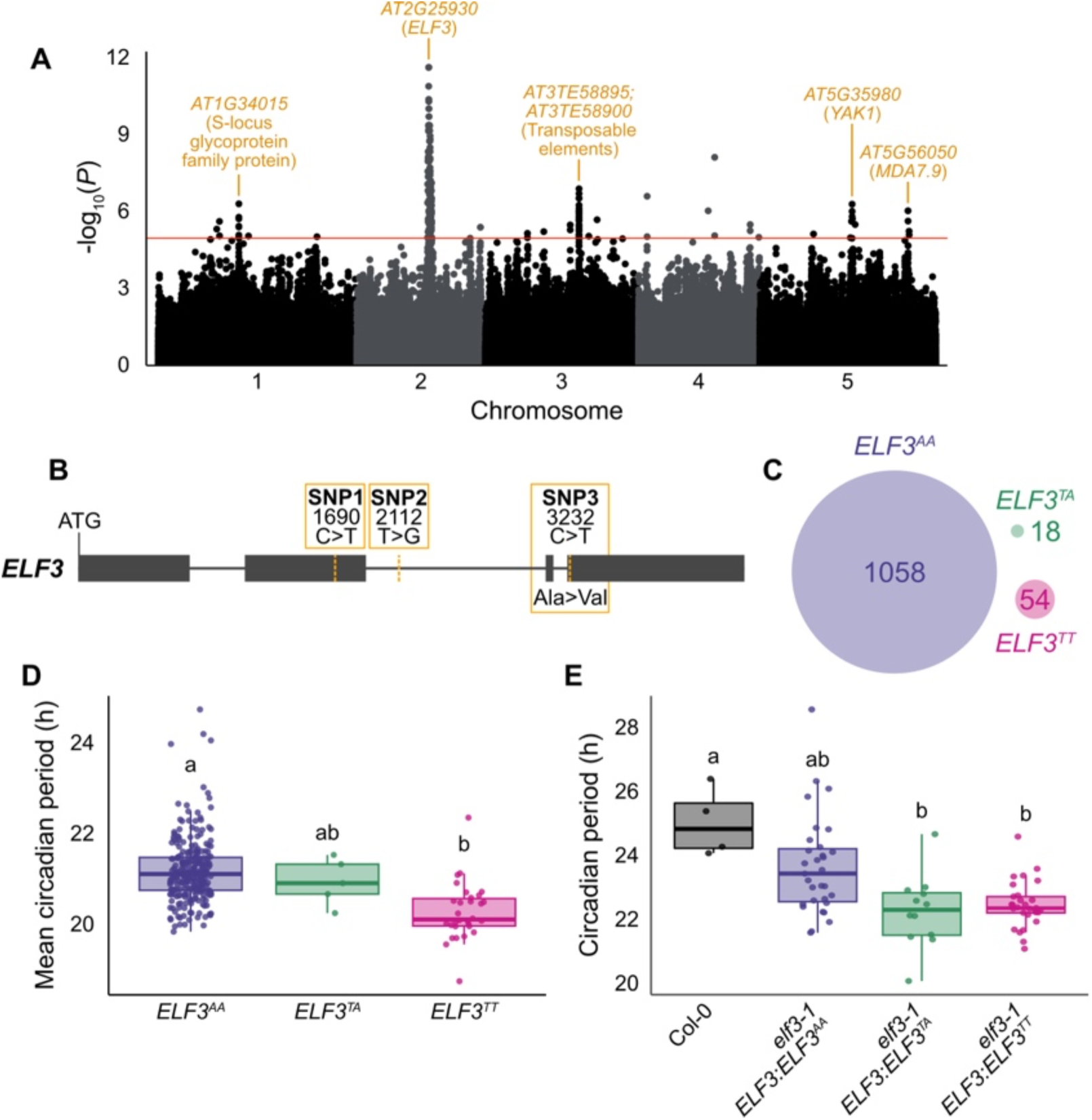
A chromosome region containing *ELF3* is significantly associated with circadian period. A, Manhattan plot summarising a genome-wide association study (GWAS) of 287 Arabidopsis accessions for circadian period. -log10(*p*) values indicate association of SNPs with circadian period. Red horizontal line is the FDR-adjusted significance threshold. Noteworthy genes within highly associated regions are annotated. B, Significantly associated SNPs within the genomic sequence of *ELF3*. C, Representation of 3 *ELF3* <u>haplogroups</u> within the 1001 Genomes collection of Arabidopsis accessions. *ELF3^AA^* accessions have Col-0 reference alleles for SNP1, SNP2 and SNP3. *ELF3^TA^* accessions have variant alleles for SNP1 and SNP2. *ELF3^TT^*accessions have variant alleles for all three SNPs. D, Distributions of mean circadian period values of Arabidopsis accessions according to *ELF3* haplogroup. E, Circadian period values of transgenic *elf3-1 ELF3p:ELF3* alleles from the three haplotypes. Haplotype groupings represent multiple independent transgenic lines (*ELF3^AA^*: 8 lines, *ELF3^TA^*: 3 lines, *ELF3^TT^*: 8 lines), n=4 for each line. Boxes in D and E represent 2^nd^ and 3^rd^ quartiles, whiskers indicate 1^st^ and 4^th^ quartiles, thick line represents the median, all data points are shown. Letters indicate significant differences as determined by one-way ANOVA followed by Tukey’s HSD; *P* < 0.05.

The most strongly associated locus was located on chromosome 2 and spanned 277 SNPs across a 480 kbp region (Figure 2A). The SNPs were in tight linkage disequilibrium, with average pairwise D’ = 0.84, which suggests that most variants are haplotype-tagging SNPs rather than causal SNPs. The most highly associated SNPs in this region (*P* < 1x10^-9^) fell within a 50 kbp sequence that contained 13 protein coding genes (Supplementary Table S3) including *ELF3*, a component of the EC and an important circadian oscillator gene. Null mutants in *ELF3* are arrhythmic (Hicks et al., 1996), but a hypomorphic *elf3-12* allele confers a short-period phenotype (Kolmos et al., 2011).

To identify which of the SNPs lying within the *ELF3* region independently explained the most variation, partial R^2^ was computed for the 277 associated SNPs on chromosome 2. The SNP that explained the highest proportion of variance (10.19%) was located in an intergenic region approximately 3 kbp downstream of *ELF3* (Chr2: 11066645). To focus on SNPs within *ELF3*, partial R^2^ analysis was repeated to include this intergenic SNP, three SNPs within the *ELF3* genomic sequence (Figure 2B) and three others within 500 bp of the genomic sequence (Chr2: 11058944-11063324) (Supplementary Table S4). In this context, the proportion of variance explained by the intergenic SNP was only 0.4%, supporting the probability that it tags variation within *ELF3* rather than being directly causal (Supplementary Table S4). Only SNPs within the *ELF3* genomic sequence had partial R^2^ values >0 (Supplementary Table S4).

There are three significantly associated SNPs within the *ELF3* genomic sequence (hereafter SNP1-3; Figure 2B). SNP1 (Chr2: 11060633), is a synonymous substitution in exon 2 of the *ELF3* transcript in a site that is poorly conserved across plant species (Supplementary Figure S4A). SNP2 (Chr2: 11061055) occurs within a relatively well conserved sequence in the second intron of *ELF3* (Supplementary Figure S4B). SNP3 (2: 11062175) is a non-synonymous polymorphism and causes an alanine to valine (A362V) substitution in a highly conserved region of the ELF3 protein sequence (Supplementary Figure S4C). SNP3 has been previously identified from QTL mapping in recombinant inbred lines and shown to contribute to variation in circadian period in the Afghan accession Shahdara (*ELF3-Sha*) (Anwer et al., 2014). The prominence of this SNP in the GWAS corroborates its importance and extends its significance to the whole Eurasian range of Arabidopsis. SNP1 and SNP2 have not been identified from previous association studies.

### A non-coding SNP in ELF3 contributes to variation in circadian period

To better understand the population structure of *ELF3,* we defined three haplogroups by combining the genotype information from SNP1, SNP2 and SNP3 across the 1001 Genomes accessions (Figure 2C; Supplementary Table S5). *ELF3^AA^* haplogroup accessions have Col-0 reference alleles for all three SNPs, *ELF3^TA^* accessions have variant alleles at SNP1 and SNP2 only, which are in complete linkage, and *ELF3^TT^* accessions, including *Sha,* have variant alleles for all three SNPs (Supplementary Table S5. No accession within the 1001 Genomes collection has an *ELF3^AT^* genotype. *ELF3^AA^* is the most frequent haplogroup, present in 1058 accessions. *ELF3^TA^* and *ELF3^TT^* are rare, present in 18 and 54 accessions, respectively (Figure 2C).

Within the 287 accessions used for circadian phenotyping, there are 252 *ELF3^AA^*, 5 *ELF3^TA^* and 30 *ELF3^TT^*accessions. Significant differences in period were identified between the haplogroups (Figure 2D). The mean circadian period was ∼0.9 h shorter in *ELF3^TT^* compared to *ELF3^AA^*. This is in agreement with Anwer et al. (2014), who showed that Arabidopsis plants carrying the non-synonymous SNP3 (*ELF3-Sha*) had shorter period lengths. *ELF3^TA^* is phenotypically intermediate to the other haplogroups, with a 0.25 h shorter mean period relative to *ELF3^AA^* (Figure 2D). Loss of function mutants in *ELF3* have an arrhythmic circadian phenotype (Hicks et al., 1996) but no significant differences in RAE were detected between accessions of the three haplogroups (Supplementary Figure S5).

The differences in circadian period between haplogroups could be confounded by background effects. To verify the functional effects of the natural *ELF3* alleles in a homogeneous genetic background, a genomic sequence including ∼2 kbp upstream of *ELF3* was cloned from representatives of each haplogroup and transformed into *elf3-1* mutants. Using at least three independent transgenic lines for each, we found that all natural *ELF3* alleles complemented the arrhythmic *elf3-1* phenotype (Supplementary Figure S6; Supplementary Table S6). However, mean circadian period of *elf3-1 ELF3:ELF3^TA^* and *elf3-1 ELF3:ELF3^TT^* was significantly shorter than wildtype Col-0 (∼2.7 h shorter) and ∼1.3 h shorter than *elf3-1 ELF3:ELF3^AA^* (Figure 2E; Supplementary Table S6). The similar period of *elf3-1 ELF3:ELF3^TA^* and *elf3-1 ELF3:ELF3^TT^* (Figure 2E; Supplementary Table S6) suggests that SNP1 and/or SNP2, neither of which affect the coding sequence of *ELF3*, confer an effect on circadian period independently of the non-synonymous SNP3.

### An intronic SNP in ELF3 affects expression of ELF3β

To test the effect of polymorphisms on *ELF3* expression we used previously published transcriptomic data (Kawakatsu et al., 2016), but found no significant difference in expression between haplogroups (Supplementary Figure S7). Thus, if changes in gene expression are caused by the SNPs, they are either subtle or condition-specific.

An alternative transcription start site has been identified in intron 2 of *ELF3*, and its product, ELF3β, is proposed to physically interact with the EC and disrupt complex formation (Wang et al., 2023). The alternative isoform *ELF3β* is not included in the transcript annotation and it is therefore possible that changes in *ELF3β* expression are associated with *ELF3* haplogroups. Since SNP2 is located within intron 2, corresponding to the region upstream of the transcriptional start site of the *ELF3β* transcript (Figure 3A), we hypothesised that SNP2 could affect the expression of this alternative transcript. To test this, we used RT-qPCR to measure full-length *ELF3α* and truncated *ELF3β* transcripts in transgenic *elf3-1* complementation lines for the three *ELF3* alleles (Figure 3B; Supplementary Table S7). The ratio of *ELF3β* to *ELF3α* expression between transgenic lines was lower in *elf3-1 ELF3:ELF3^TA^* and *elf3-1 ELF3:ELF3^TT^*lines relative to *elf3-1 ELF3:ELF3^AA^* lines and this difference was significant for *ELF3^TA^* (Figure 3B). Thus, the non-reference SNP2 variant might disrupt an enhancer sequence causing reduced expression of *ELF3β*. It is notable that SNP3 is not within the coding sequence of the *ELF3β* transcript (Wang et al., 2023; Figure 3A) and therefore is not expected to affect function of ELF3β. Thus, the functional contributions of SNP2 and SNP3 are likely to be independent.

**Figure 3:**
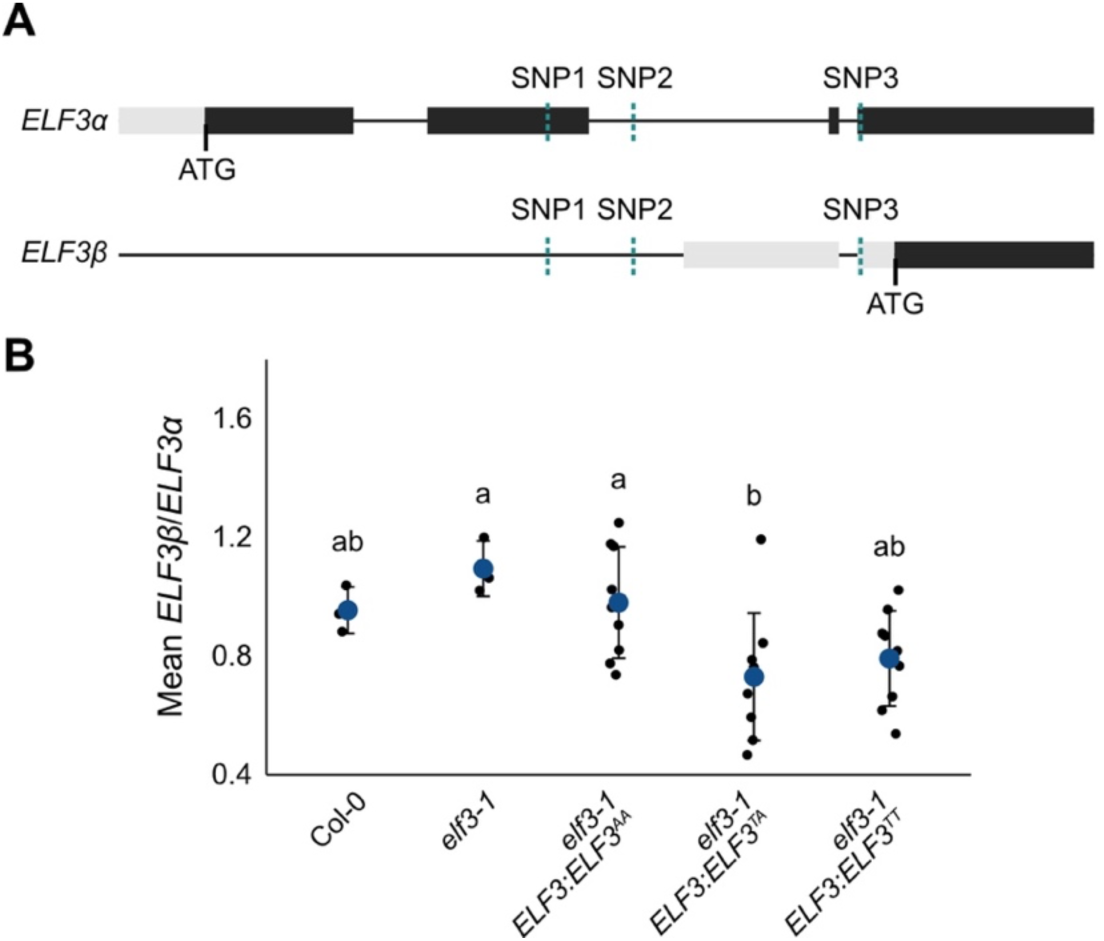
*ELF3^TA^* and *ELF3^TT^* alleles have reduced expression of *ELF3β*. A, Gene models of the main isoform of ELF3 (*ELF3α*) and the alternative truncated isoform (*ELF3β*) with the positions of SNP1-3 indicated. B, Ratio of *ELF3β* to *ELF3α* transcript level in Col-0, *elf3-1* and transgenic *elf3-1 ELF3p:ELF3* alleles from the three haplotypes. Each haplotype allele is represented by three individual transgenic lines. Blue circles are means±SD, n=3 for each line. Letters indicate significant differences as determined by one-way ANOVA followed by Tukey’s HSD; *P* < 0.05.

### Evidence of a selective sweep in ELF3 haplogroups in response to climate

Since SNP1/2 and SNP3 appear to both contribute to circadian period differences in natural Arabidopsis accessions, we searched for evidence of selection occurring at the *ELF3* locus across the Arabidopsis species range. Tajima’s *D* was computed across the genome for the different haplogroups (Figure 4A). Negative Tajima’s *D* values indicate that rare mutations are common within a given sequence, which is a signature of purifying selection (Carlson et al., 2005).

**Figure 4:**
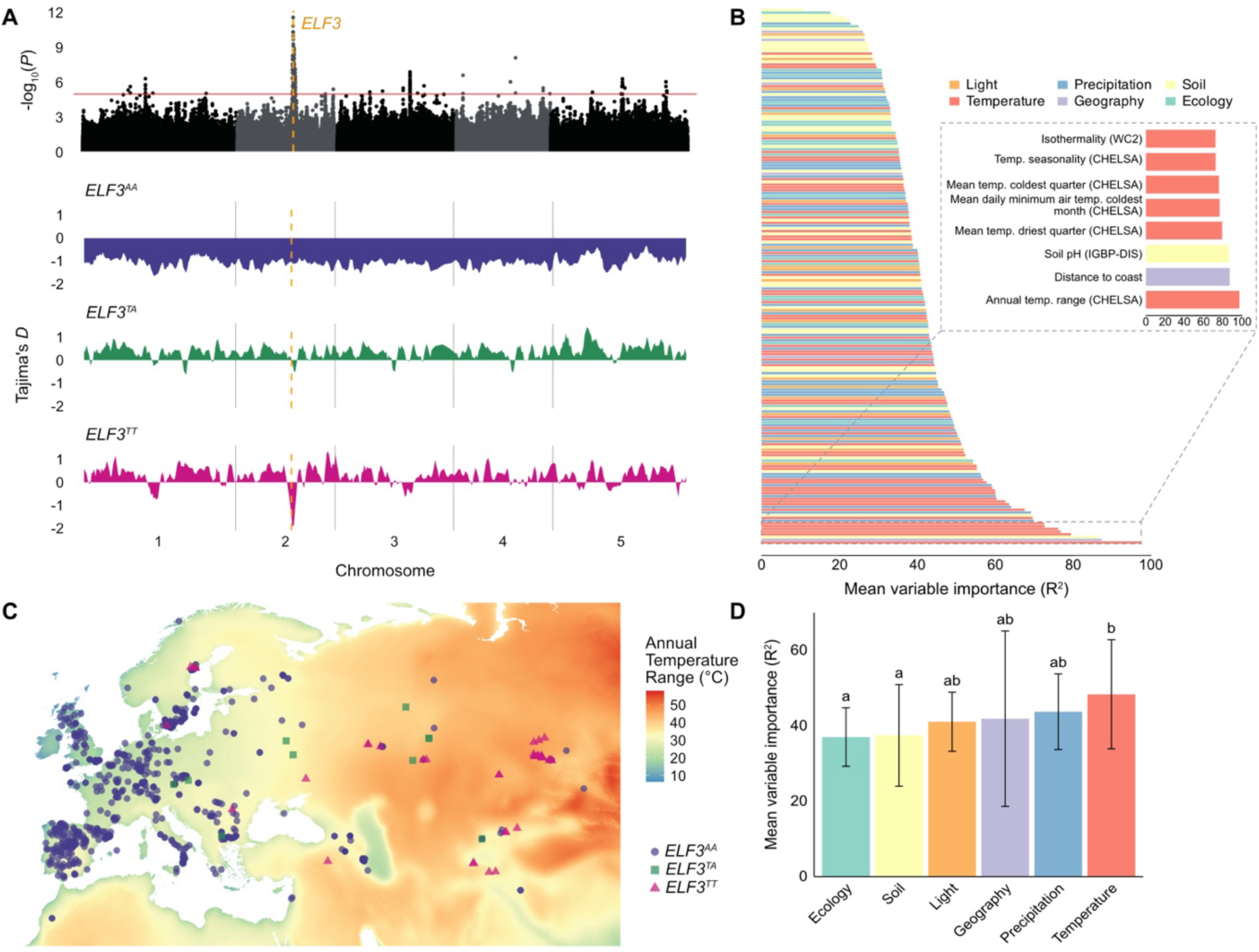
*ELF3* SNPs have been selected for in continental climates. A, Tajima’s *D* was calculated for accessions of each *ELF3* haplogroup in sliding windows of 5 kbp with 1 kbp steps across the entire genome. The genomic location of *ELF3* is indicated on the GWAS Manhattan plot and Tajima’s *D* distribution plots by a dashed orange line. B, Random forest regression of *ELF3* SNP identity against 196 bioclimatic variables. Random forest regressions were performed for the 3 SNPs individually and variable importance values were averaged. C, Distribution of the most important bioclimatic variable in B. Annual temperature range (CHELSA BIO7) reflects the geographical range of *ELF3* haplogroups across Eurasia. D, Mean variable importance values from the random forest regressions were averaged across categories of bioclimatic variables. Error bars represent ±sd Letters indicate significant differences as determined by one-way ANOVA followed by Tukey’s HSD; *P* < 0.05.

Across the entire genome, average Tajima’s *D* for *ELF3^AA^*, *ELF3^TA^* and *ELF3^TT^* accessions was -1.07, 0.38 and 0.30, respectively (Figure 4A). Within the genomic sequence of *ELF3*, *D* values for all three *ELF3* haplogroups were negative (Figure 4A), although *D* in *ELF3^AA^* (-1.75) and *ELF3^TT^* (-1.62) was much lower than in *ELF3^TA^*(-0.38). Given the context of a genome-wide trend towards negative *D* in *ELF3^AA^* accessions (Figure 4A), it is likely that the negative value at the *ELF3* locus for *ELF3^AA^* reflects a recent population expansion rather than purifying selection. On the other hand, the strongest signature of positive selection throughout the genome in *ELF3^TT^* accessions fell precisely at the *ELF3* locus, whilst *ELF3^TA^* also exhibited a negative trend in *D* at this position (Figure 4A). A null distribution of Tajima’s *D* was computed to test the significance of these values. Less than 2.5% of null values (significance threshold) were lower than the *D* values observed in *ELF3^TT^*, whereas over 50% of the null distribution was lower than *D* for *ELF3^TA^*. This evidence strongly suggests that the *ELF3* locus has undergone a selective sweep amongst *ELF3^TT^* accessions and, to a lesser extent, *ELF3^TA^* accessions.

The three *ELF3* haplogroups are found in geographically distinct locations; the large *ELF3^AA^* group primarily occupies Western Europe, *ELF3^TT^* is located in Eastern Europe and Central Asia, and *ELF3^TA^* is geographically intermediate (Figure 4C). To determine what might be driving the distribution of haplogroups, random forest was used to classify the haplogroups based on 196 bioclimatic variables at the location of origin of the accessions (Fournier-Level et al., 2022). Variables related to temperature explained the presence of SNP alleles better than other categories (Figure 4B-D). When averaged across the three SNPs, the most important variable was annual temperature range (Figure 4B,C; Supplementary Table S8). In particular, the magnitude of annual temperature range, which is indicative of continental climate, broadly matches the distribution of *ELF3* haplogroups across the landscape (Figure 4C).

To infer evolutionary relationships between haplogroups, a maximum likelihood (ML) phylogenetic tree was constructed from *ELF3* sequences of the three haplogroups (Figure 5A). All accessions from *ELF3^TT^* and *ELF3^TA^*were grouped into a monophyletic clade that diverged from *ELF3^AA^*(Figure 5A). This suggests that the *ELF3^TA^* haplogroup was not a stepping stone in the evolution of *ELF3^TT^*. The *ELF3^TT^* haplogroup formed two distinct clades within the tree, one with a single accession Kz-13 and the other comprising all remaining *ELF3^TT^*accessions from which the *ELF3^TA^* haplogroup diverged (Figure 5A). Thus, it is possible that *ELF3^TA^* haplogroups arose through reversion from *ELF3^TT^*. However, a previous study identified evidence for admixture of sequences from ‘relict’ (i.e. pre-glacial) populations in the Kz-13 genome (Alonso-Blanco et al., 2016), which could explain its dissimilarity to other *ELF3^TT^* accessions. A time-calibrated coalescence analysis was used to estimate divergence times between haplogroups. The consensus analysis places *ELF3^AA^*as the ancestral haplogroup, with mean time to most recent common ancestor (tMRCA) 7.5 kya (Figure 5B). Consistent with evolutionary relationships in the ML tree, *ELF3^TA^*and *ELF3^TT^* diverged later (4.3 kya and 4.5 kya, respectively) (Figure 5B).

**Figure 5:**
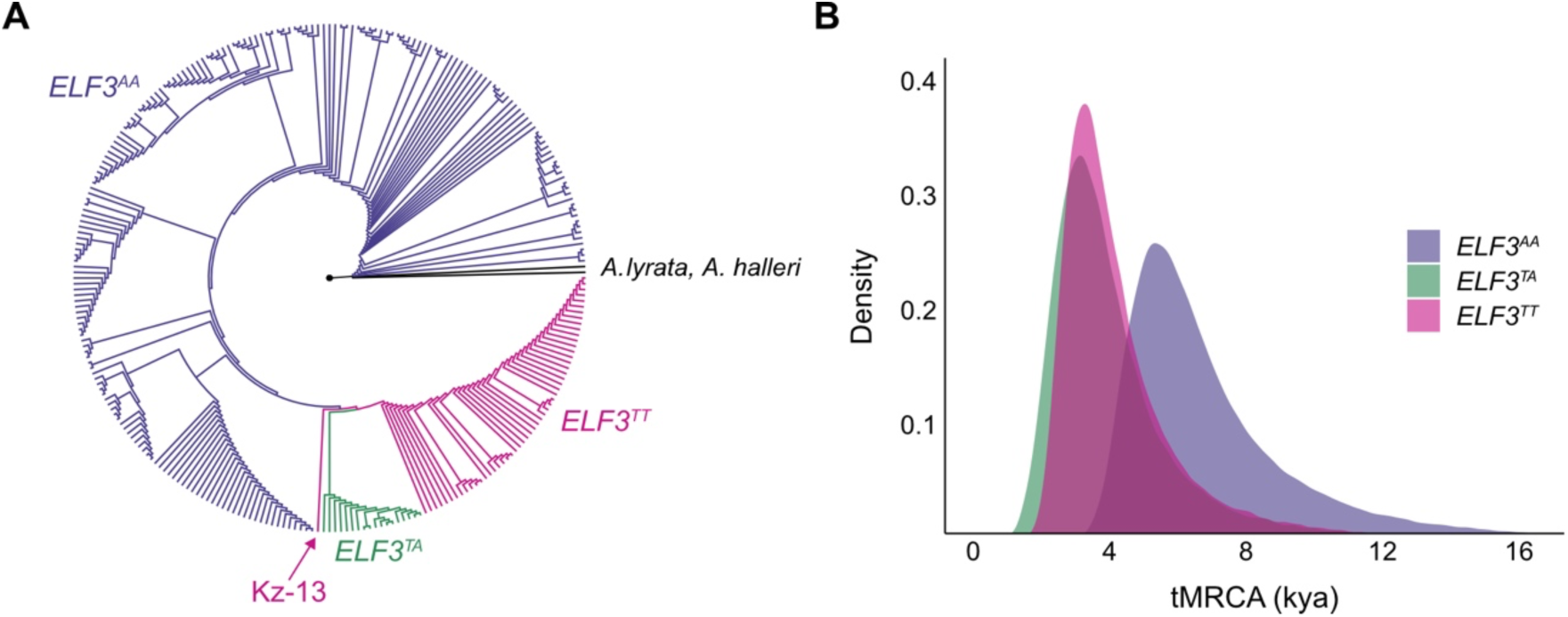
The evolutionary history of *ELF3* haplogroups. A, Maximum likelihood tree of *ELF3* sequences from each haplogroup determine by phylogenetic analysis. The sequences of *Arabidopsis lyrata* and *Arabidopsis halleri* were used as outgroups. B, Bayesian coalescence analysis of divergence times between *ELF3* haplogroups. Density of estimates of tMRCA across 1.2x10^8^ MCMC generations are plotted.

The combination of geographical distribution and evolutionary relationship of the haplogroups suggest that the mutations in *ELF3* might have contributed to effective responses to temperature in highly variable, continental climates. Indeed, the *ELF3-Sha* allele, which belongs to the *ELF3^TT^* haplogroup, underlies a QTL for high-temperature hypocotyl elongation (Raschke et al., 2015). To assess the contribution of *ELF3* polymorphisms to the temperature response, hypocotyl elongation of five accessions representing each of the three *ELF3* haplogroups was tested under different temperature regimes: constant temperature (28°C or 22°C) or daily temperature cycles (28°C/12°C or 22°C/7°C) (Supplementary Table S9). In constant 22°C or 22°C/7°C cycles, there were no significant differences in hypocotyl length between haplogroups (Figure 6A; Supplementary Figure S8A). In 28°C/12°C cycles, the hypocotyl length of the haplogroups or *elf3-1* mutants did not differ (Figure 6B). However, there were large significant differences between the hypocotyl lengths of haplogroups in constant 28°C (Figure 6B; Supplementary Figure S8B). Interestingly, the average hypocotyl length of each haplogroup in warm temperature, as well as the fold change in hypocotyl length between constant 22°C and constant 28°C (Figure 6C) reflected the same trend in differences between haplogroups for circadian period (Figure 2D). That is, the response of *ELF3^TA^* accessions was intermediate to the responses of *ELF3^AA^* and *ELF3^TT^* accessions (Figure 6C). We also tested whether *ELF3* polymorphisms affect the response of flowering at elevated temperature. While *ELF3^TA^* and *ELF3^TT^* accessions flowered later than *ELF3^AA^* accessions, the decrease in time to flowering in response to elevated temperature was relatively larger in these haplogroups (Supplementary Figure S9; Supplementary Table S10). Together, these results suggest a functional contribution of the *ELF3^TA^* haplogroup to temperature responses, as well as circadian period.

**Figure 6:**
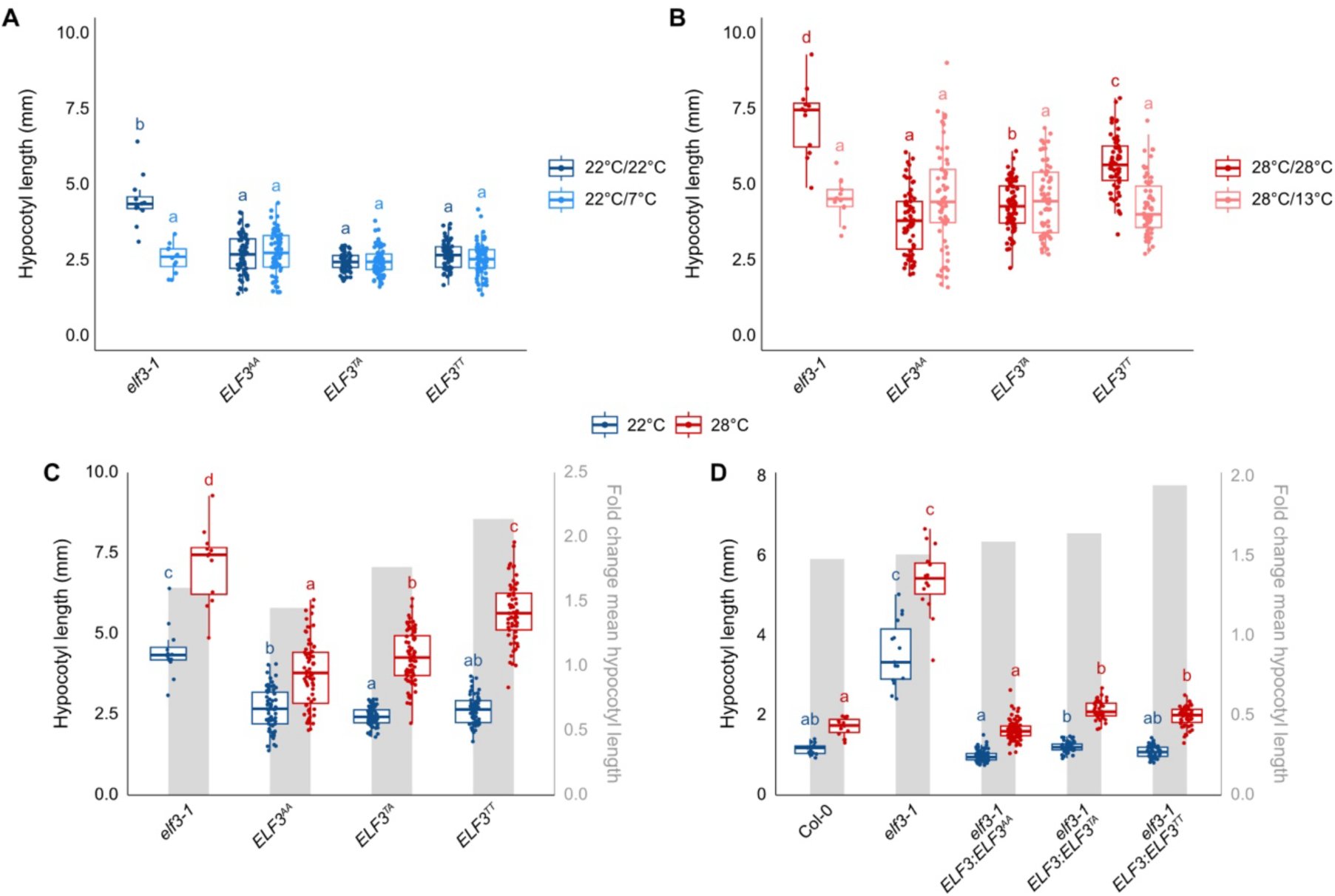
Naturally occurring SNPs in *ELF3* are associated with enhanced thermomorphogenesis. A-B, Hypocotyl length of five accessions representing each *ELF3* haplogroup (*ELF3^AA^*: Col-0, IP-Lac-0, Zu-0, Ull2-3, Cvi-0; *ELF3^TA^*: Sku-30, Fäb-2, Uod-1, Per-1, Sij-4; *ELF3^TT^*: Omn-5, Kolyv-2, Lebja-1, Noveg-3 and Shahdara) and *elf3-1* mutants grown constant control temperature (22°C/22°C), cycling control temperature (22°C/7°C), constant elevated temperature (28°C/28°C) or cycling elevated temperature (28°C/12°C). C, Hypocotyl length (left axis) and fold change of mean hypocotyl length (grey boxes, right axis) between constant 22°C and 28°C for *elf3-1* and natural accessions in A and B, n=16. D, Hypocotyl length (left axis) and fold change of hypocotyl length (grey boxes, right axis) between constant 22°C and 28°C for transgenic *elf3-1 ELF3p:ELF3* alleles from the three haplotypes. Haplotype groupings represent multiple, independent transgenic lines (*ELF3^AA^*: 9 lines, *ELF3^TA^*: 5 lines, *ELF3^TT^*: 6 lines), n=16 for each line. Letters indicate significant differences within each temperature condition as determined by one-way ANOVA followed by Tukey’s HSD; *P* < 0.05.

To test the effect of the three *ELF3* alleles in a homogeneous genetic background we measured hypocotyl length in the *ELF3* transgenic lines in constant 22°C or 28°C (Figure 6D; Supplementary Table S11). We found significantly longer hypocotyl for *ELF3^TA^*and *ELF3^TT^* transgenics in warm temperature than *ELF3^AA^*transgenics or Col-0, and a temperature response in *ELF3^TA^* that was intermediate to the *ELF3^AA^* and *ELF3^TA^* transgenics. Together, these results suggest that the non-coding SNP1 and/or SNP2 in *ELF3* confer a functional effect on temperature responses of Arabidopsis accessions that might have been selected for in natural populations.

## Discussion

We have quantified the extent of variation in circadian rhythms in natural accessions representing the geographical range of Arabidopsis and found substantial, heritable variation in circadian period across 287 accessions. GWAS of circadian period revealed that extant natural variation of circadian period in these accessions is largely modulated by *ELF3*. Variation in *ELF3* has been selected for and might have aided the spread of Arabidopsis towards Eastern Europe and Central Asia. In keeping with previous studies that have identified thermosensory functions of ELF3, we found evidence that the forces driving the spread of *ELF3* variants are related to elevated temperatures.

Three SNPs in *ELF3* were identified as putative candidates for the adjustment of circadian period in natural accessions. The non-synonymous SNP (SNP3) shared by *ELF3^TT^* accessions has previously been identified in the Bay-0 x Sha mapping population (Anwer et al., 2014). The *ELF3-Sha* allele was implicated in the regulation of circadian period (Anwer et al., 2014), as well as temperature-induced hypocotyl elongation (Raschke et al., 2015), and the causal effect of the A362V peptide change induced by SNP3 was confirmed in both cases, and attributed to improper localisation of the variant protein to the nucleus (Anwer et al., 2014). Our data supports a contribution of this *ELF3* genotype to natural variation in circadian period and provides evidence of selection in natural populations.

The effects or functions of SNP1 and SNP2 have not been previously investigated. SNP1 causes a synonymous change in exon 2 of the *ELF3* gene sequence. Synonymous SNPs can affect processes such as transcription factor binding, mRNA stability, and translation elongation (Gustafsson et al., 2004; Agashe et al., 2013; Zwart et al., 2018; Shen et al., 2022). SNP1 converts the underrepresented Gly-encoding GGC codon to a more optimal GGU codon (Duret and Mouchiroud, 1999), which could potentially affect translation efficiency or protein folding. The potential functional effects of intronic SNPs are often overlooked. An intronic SNP within the barley (*Hordeum vulgare*) *ELF3* sequence causes intron retention and a putative truncated protein (Xia et al., 2017). SNP2 in intron 2 of *ELF3* is upstream of a transcriptional start site for the *ELF3β* transcript (Wang et al., 2023) and our data suggests that SNP2 affects expression of this alternative transcript (Figure 3). ELF3β is proposed to repress the activity of the EC. Interestingly, the *ELF3β* promoter includes putative heat shock factor (HSF) binding motifs (Wang et al., 2023). While the synonymous SNP1 falls within exon 2 of *ELF3α*, it is also positioned upstream of the *ELF3β* TSS (Figure 3A). Thus, SNP1 and SNP2 could affect temperature-dependent expression of *ELF3β* in *ELF3^TA^* and *ELF3^TT^* haplogroups. It is intriguing that functional variation in the promoter of *ELF3β* has arisen in Arabidopsis as well as pear (Wang et al., 2023), with effects on flowering and temperature-responsive growth. *ELF3β* might therefore have strong potential as a breeding target for maturation and temperature resilience in crops. More widely, these results highlight the need for the consideration of non-coding genetic variation in research and breeding programs.

Evolutionary studies of Arabidopsis suggest that extant Eurasian populations are almost exclusively descendants of a post-glacial expansion of ‘non-relicts’ that replaced relict populations of ice age refugia from 10 kya (François et al., 2008; Alonso-Blanco et al., 2016; Lee et al., 2017; Hsu et al., 2019; Toledo et al., 2020). The divergence of the *ELF3* haplogroups was predicted to have occurred around 7.5 kya (Figure 5B), consistent with this non-relict expansion. Our phylogeny and coalescence modelling supported the ancestral nature of the *ELF3^AA^* haplogroup, while Eastern European and Central Asian *ELF3^TA^* and *ELF3^TT^* haplogroups diverged later (Figure 5). Given the contemporary range of haplogroups (Figure 4C), this suggests that the geographical direction of gene flow and selection within *ELF3* has occurred from west to east. This is consistent with previous work, which suggested that non-relict populations expanded in westward and eastward directions from a Balkan or Black Sea origin (Lee et al., 2017). It is noteworthy that there are admixed accessions in the *ELF3^TA^*and *ELF3^TT^* haplogroups (Chi-0 from *ELF3^TA^*, Kz-13 and Nemrut-1 from *ELF3^TT^*). Alonso-Blanco et al. (2016) found evidence for historic introgression of genetic information from relict Mediterranean populations among admixed accessions. Given that Kz-13 represented a discrete, more ancestral clade of *ELF3^TT^* in the ML tree (Figure 5A), this could indicate that haplogroup divergence occurred in regions where relicts would have been encountered (e.g. Mediterranean refugia). Together, our demographic and evolutionary analyses suggest that variation in *ELF3* was important for the historic spread of Arabidopsis into environments with novel temperature demands.

Our finding of historic selection of natural variation in *ELF3* in Arabidopsis builds on the emerging story of wide-ranging importance of *ELF3* to the success of plants. *ELF3* variants in Arabidopsis alter numerous phenological and physiological traits, including flowering time (Reed et al., 2000), senescence (Kim et al., 2018) and temperature responsiveness (Raschke et al., 2015). An association study of flowering time in populations of ragweed (*Ambrosia artemisiifolia*) identified *ELF3* as a primary candidate (Battlay et al., 2023). As ragweed became established in northern Germany, the allele frequency of a flowering time-associated non-synonymous SNP in *ELF3* increased 28-fold between historic and modern ragweed populations (Battlay et al., 2023). *ELF3* is also a prominent marker in GWAS of crop cultivars. *ELF3* was associated with multiple agricultural and developmental traits in a diverse barley population (Maurer et al., 2016), and an association of *ELF3* with flowering time has been reported in the tropical legume pigeonpea (*Cajanus cajan*; Varshney et al., 2017). Variation in *ELF3-D1* in bread wheat (*Triticum aestivum*) affects flowering time, maturation and grain nutrition (Wittern et al., 2023; Buckley et al., 2025). The effect of ELF3 on phenology in barley and wheat interacts with temperature (Ochagavía et al., 2019; Zhu et al., 2023), which further suggests that *ELF3* might be an ideal target for modulation of temperature plasticity in future climate-proof crops.

Polymorphism in *ELF3* aided the eastward spread of Arabidopsis populations towards continental climates, with a particularly strong signature of selection identified among *ELF3^TT^*accessions. In the absence of a latitudinal cline in circadian period, other abiotic factors such as temperature variability might have exerted a stronger selection pressure on clock genes. The phenotypic effects of SNPs in *ELF3* on circadian rhythms and temperature responses highlight the multifaceted roles of clock proteins, which exert influence on many aspects of plant biology through their hundreds or thousands of genomic targets (Haydon et al., 2019). However, this could also pose challenges for the reengineering of crop development through ELF3, since existing pleiotropic constraints could result in maladaptation of *ELF3* variants. To avoid this, breeders could co-opt naturally occurring *ELF3* polymorphisms that might have more subtle functional effects. This study details one example of how altered clock function has been used for adaptation to different temperature regimes, and there are likely other examples of clock gene selection for niche adaptation in natural plant populations.

## Materials and methods

### Plant materials and growth conditions

A collection of 287 *A. thaliana* accessions from the 1001 Genomes Project (Alonso-Blanco et al., 2016) was used for phenotyping experiments (Supplementary Table S12). Accessions were selected to capture diversity for nine bioclimatic variables at their site of origin retrieved from the AraCLIM database (latitude and longitude, average temperature and temperature range, solar insolation, net radiation, normalised difference vegetation index (NDVI), cloud cover, UV index; Ferrero-Serrano and Assmann, 2019). Col-0 and the circadian clock gene mutants *cca1-1* (Green and Tobin, 1999) and *ztl-105* (Martin-Tryon et al., 2007) were used as controls.

Unless stated otherwise, Arabidopsis plants were grown using the following standard protocol. Arabidopsis seeds were surface sterilised and sown on half-strength Murashige and Skoog (½ MS) media (0.8 % agar type M; Sigma-Aldrich). Seeds were chilled at 4°C in the dark for 48 h before transfer to a HiPoint 740FHLED growth chamber (HiPoint Corporation) set to 20°C under a 12 h light/ 12 h dark (LD) cycle with approximately 100 µmol m^−2^ s^−1^ photosynthetically active radiation (PAR).

### Transient luciferase seedling transformation

The transient luciferase protocol was conducted following Ting et al. (2022) with modifications (Supplementary Figure S1). Seeds were sown on ½ MS, 1% (w/v) agar in 60 mm petri dishes and chilled for seven days before transfer to a growth cabinet. *A. tumefaciens* (C58) carrying a *GIGANTEA* (*GI*) promoter-driven luciferase reporter construct (*pE301-GIp:LUC2*) were inoculated into liquid LB media and incubated at 30°C. After 24 h, these were subcultured into a 10-fold larger volume of LB-MES media (LB buffered with 1 M MES to a final pH of 5.5 with KOH) with 200 µM acetosyringone and incubated for a further 18 h. The culture was resuspended in an equal volume of infiltration buffer (50 mM MES, 10 mM MgCl_2_, 200 µM acetosyringone, pH 5.5). Transformations were performed with 5 d old seedlings at *Zeitgeber* Time 4 (ZT4). Seedlings were submerged in approximately 10 mL of culture and infiltrated 2x 60 s in a vacuum desiccator. Five minutes after vacuum infiltration, the culture was discarded and plates were returned to the growth cabinet. Six d-old seedlings were sprayed with 1 mM D-luciferin K^+^ salt at ZT0 using a sterile atomiser (approximately two sprays per plate). After 24 h, clusters of ten transformed seedlings were transferred to black 96-well plates containing 300 µL ½ MS media (0.8% agar, 50 µg/mL timentin) per well. Approximately 25 µL 1 mM D-luciferin K^+^ salt was applied to each seedling cluster before plates were placed into an imaging box. Seedlings were maintained in LD conditions until subjective dawn (ZT0) and luciferase luminescence was measured hourly for 120 h in continuous light (LL). A total of nine batches were required for phenotyping of the 287-accession panel, with four individual replicates of accessions or controls in each batch.

Luciferase luminescence was imaged using a Retiga LUMO charge-coupled device (CCD) camera (Teledyne Photometrics) fitted with a 25 mm f/0.95 lens housed within a light-tight dark box. Light (>45 µmol m^-2^s^-1^) was provided by two LB3 red/blue LED panels (Photek). The lights and camera were controlled by beanshell (bsh) scripts in µManager (version 2.0; Edelstein et al., 2014). Luciferase luminescence was measured by 10 min camera exposure in the dark, following 45 min of light and 5 min in the dark to exclude capture of delayed leaf fluorescence (Gould et al 2009). Camera properties were binning = 4x4, gain = 1, readout rate = 0.650195 MHz 16 bit. Luminescence was measured in seedling clusters from image stacks as integrated density using ImageJ (NIH). Timeseries data were uploaded to BioDare2 (Zielinski et al., 2014) for period analysis of background-subtracted data using the fast Fourier transform non-linear least squares (FFT*-*NLLS) method (Plautz et al., 1997). Data were detrended according to the amplitude and baseline detrending method and analysis was performed using a 24 - 120 h window. Estimates of period that fell outside of the range of 18 - 30 h were excluded. Time series intensity data presented are Z-score normalised.

### Temperature-dependent hypocotyl elongation

For each genotype, 16 seeds were sown onto 10cm^2^ Petri plates with ½ MS media, 0.8% (w/v) agar. Plates were positioned vertically within growth cabinets and exposed to LD cycles and germinated at 22°C for two days. Plates were then transferred to one of four 12 h/12 h temperature regimes; 22°C/7°C, 22°C/22°C, 28°C/13°C or 28°C/28°C. After 5 days of treatment, plates were imaged and hypocotyl length was quantified using ImageJ (Schindelin et al., 2012).

### Temperature-dependent flowering time

Six replicates of 5 accessions from each of the three haplogroups and *elf3-1* were sown directly to soil (Australian Growing Solutions Plugger seed raising mix with 1.5 % v/v perlite) and germinated in 12: 12 h LD cycles at 22°C. Trays of 21 day-old plants were vernalised for 28 days in 12°C: 4°C cycles and then transferred to either 22°C or 28°C constant temperature. At bolting, the number of rosette leaves from each plant was counted to indicate the developmental age at flowering.

### Genome-wide association study

Period, phase, amplitude and relative amplitude error (RAE) were analysed for each of the 287 accessions using linear-mixed models (LMM) implemented in the R\lme4 package (Bates et al., 2015). Batch and accession were modelled as random effects, and variance among accessions was used to compute broad-sense heritabilities. Best linear unbiased predictors (BLUPs) of the accession effect for each trait were used for genome-wide association studies (GWAS) with genotype data procured from the 1001 Genomes data centre (https://1001genomes.org/data-center.html). Genome wide association studies (GWAS) were conducted using a LMM accounting for the random effect of population structure through the inclusion of a IBS kinship matrix (computed using PLINK 2.0; Chang et al., 2015) and testing the fixed effect of single nucleotide polymorphisms (SNPs) as implemented in GEMMA (v0.98.5; Zhou and Stephens, 2012). SNPs with a minor allele frequency below 0.05 were filtered out and *p*-values obtained from a Wald test of the SNP effects were adjusted for a false discovery rate FDR = 5% (Benjamini and Hochberg, 1995). Partial coefficients of determination (partial R^2^) were calculated for significantly associated SNPs. Linkage disequilibrium between SNPs was estimated using pairwise D’ as implemented in PLINK (Chang et al., 2015).

### Random forest regression of bioclimatic variables

The allele turnover over environmental gradients for associated SNPs within *ELF3* was inferred through random forest regression (Fournier-Level et al., 2022) for 196 quantitative environmental variables retrieved from the AraClim database (Ferrero-Serrano and Assmann, 2019) using the R\randomForest package (Liaw and Wiener, 2002). The number of variables randomly sampled for each tree was set to the square root of the total number of variables (√196= 14); the number of trees to grow was set to 500. Variable importance of each environmental variable in the final model was computed as the percentage of decrease in accuracy when the variable was excluded. Data presented display the mean variable importance values for three individual SNPs.

### Multiple sequence alignment

To assess evolutionary conservation of SNPs, orthologous *ELF3* sequences from maize (*Zea mays*), rice (*Oryza sativa*), barley (*Hordeum vulgare*), wheat (*Triticum aestivum*), barrel clover (*Medicago truncatula)*, tomato, soybean and rapeseed (*Brassica napus*) were identified through BLAST. Gene sequences were aligned using the MUSCLE (Edgar, 2004) algorithm implemented in Mega 11 (Tamura et al., 2021) with default settings.

### Generation of ELF3 transgenic lines

The genomic *ELF3* sequence, including ∼2000 bp upstream from the start codon to the stop codon were amplified by PCR from gDNA from Col-0 (*ELF3^AA^*), Sij-4 (*ELF3^TA^*) and Lebja-1 (*ELF3^TT^*) using Phusion DNA polymerase (NEB) and primers listed in Supplementary Table S13. A-tailed products were inserted into the pCR8/GW/TOPO entry vector (Invitrogen) using a TOPO reaction and introduced into *pEarleyGate301* (Earley et al., 2006) using LR Clonase II (Invitrogen). *ELF3* sequences were confirmed by Sanger sequencing (Supplementary Table S13). Constructs were introduced into *Agrobacterium tumefaciens* C58 and transformed into *elf3-1* plants (Col-0 background) by floral dip (Clough and Bent, 1998). Phosphinothricin (PPT)-resistant T2 or T3 plants were used for phenotyping experiments. For each *ELF3* variant, at least three independently transformed transgenic lines were used in each experiment.

### Analysis of selective sweeps

To test for deviation from neutrality, Tajima’s *D* was computed within each *ELF3* haplogroup for the 345 SNPs located within *ELF3* using the R\snpR package (Hemstrom and Jones, 2023). The most frequent haplogroup - *ELF3^AA^*, present in 1058 - accessions was randomly downsampled to 200 accessions. The Spanish accession IP-Cad-0 was excluded from further analysis due to a suspected genotyping error within *ELF3* since it is an outlier relative to other *ELF3^TA^* accessions with respect to its location of origin. To assess significance, a null distribution for Tajima’s *D* for each haplogroup was drawn through 1000 permutations of random assignation of accessions to haplogroups. Observed *D* values that exceeded the two-tailed 5th percentile of the null distribution were deemed significant. Measures of Tajima’s *D* confound selection processes affecting specific loci with demographic processes affecting the whole genome. To distinguish between these, a genome-wide mean Tajima’s *D* was calculated across sliding windows of 5 kbp, moving 1 kbp each step.

### Phylogenetic and evolutionary divergence analysis

*ELF3* haplogroup sequences were downloaded from the 1001 Genomes Tools site (https://tools.1001genomes.org/pseudogenomes/) with the same downsampling of 200 *ELF3^AA^* accessions described above. *ELF3* genomic sequences for *Capsella rubella* and *Camelina sativa* were included as an outgroup to the *Arabidopsis* genus, and *Arabidopsis lyrata* and *Arabidopsis halleri* sequences were included as close relatives of *A. thaliana*. Sequences were aligned according to the method described above. The most suitable substitution model for the haplogroup sequences, HKY+G (Hasegawa et al., 1985) was determined using jModelTest 2 (Darriba et al., 2012). A maximum likelihood (ML) tree was constructed using PhyML 3.0 (Guindon et al., 2010). Branch supports were calculated by approximate likelihood ratio test (aLRT SH-like). The ML tree was visualised as a cladogram with FigTree v1.4.4 (http://tree.bio.ed.ac.uk/software/figtree/).

Divergence time of *ELF3* sequences between haplogroups was estimated using a Bayesian coalescent model implemented in BEAST v2.7.5 (Bouckaert et al., 2019). The calibrated Yule model was used as the tree prior (Heled and Drummond, 2012), and three calibration points were set based on previous estimations (Hohmann et al., 2015; Hsu et al., 2019). The divergence time of *Capsella*, *Camelina* and *Arabidopsis* (i.e. the tree root) was set to 8.16 mya, the divergence of *Capsella* and *Camelina* at 7.36 mya and the divergence time within the *Arabidopsis* genus was set to 5.96 mya. Divergence times were estimated using normal prior distribution with the above means and a standard deviation of 1 mya for all priors. A log-normal, uncorrelated relaxed clock was used as the molecular clock (Drummond et al., 2006; Douglas et al., 2021), with starting rate 7x10^3^ substitutions/site/million years based on the estimated mutation rate for *A. thaliana* (Ossowski et al., 2010). An uninformative uniform prior was used for the molecular clock rate to avoid narrowing the search space (Bromham et al., 2018). Three Markov chain Monte Carlo (MCMC) simulations were run for 5x10^7^ iterations with a burn-in period of 1x10^7^, giving a final combined chain length of 1.2x10^8^. Chains were combined using LogCombiner v1.10.4 (Drummond and Rambaut, 2007), and Tracer v1.7.2 (Rambaut et al., 2018) was used to visualise convergence based on effective sample size (ESS) for each chain.

### RT-qPCR

Arabidopsis seeds of *elf3-1 ELF3p:ELF3* transgenic lines, Col-0 and *elf3-1* were grown at 22°C for ten days. Ten-seedling clusters were harvested in triplicate immediately before dusk (ZT12) and flash frozen in liquid nitrogen prior to storage at -80°C until processing. RNA was extracted from ground frozen tissue using ISOLATE II RNA Plant Kit (Meridian Bioscience). cDNA was prepared from 0.5 µg RNA in 10 µl reactions using Tetro cDNA synthesis kit (Bioline). 5 ng/µl of cDNA was used in each PCR reaction with 0.4 µM primers in the SensiFAST SYBR no-ROX kit (Bioline) on a CFX Opus 384 Real-time PCR system (BioRad). PCR reaction efficiencies were determined for each primer pair using LinRegPCR (Ruijter et al., 2009) and transcript levels were determined for target and reference genes (UBQ10) using (mean PCR efficiency)^-Ct^. Primer sequences are listed in Supplementary Table S13.

## Acknowledgements

Research was funded by the Research Continuity Scheme from the Faculty of Science, University of Melbourne to MJH and a Melbourne Research Scholarship and Faculty of Science Postgraduate Writing-Up Award to CRB.

## Author contributions

MJH conceived the study, CRB, AFL and MJH designed experiments, CRB and MJH acquired data, CRB, AFL and MJH analysed and interpreted data, CRB drafted the manuscript and prepared figures, CRB, AFL and MJH reviewed and revised the manuscript.

