## Supplemental Figures for "A non-coding SNP in *ELF3* alters expression of *ELF3β* and confers adaptation of Arabidopsis to a continental climate"

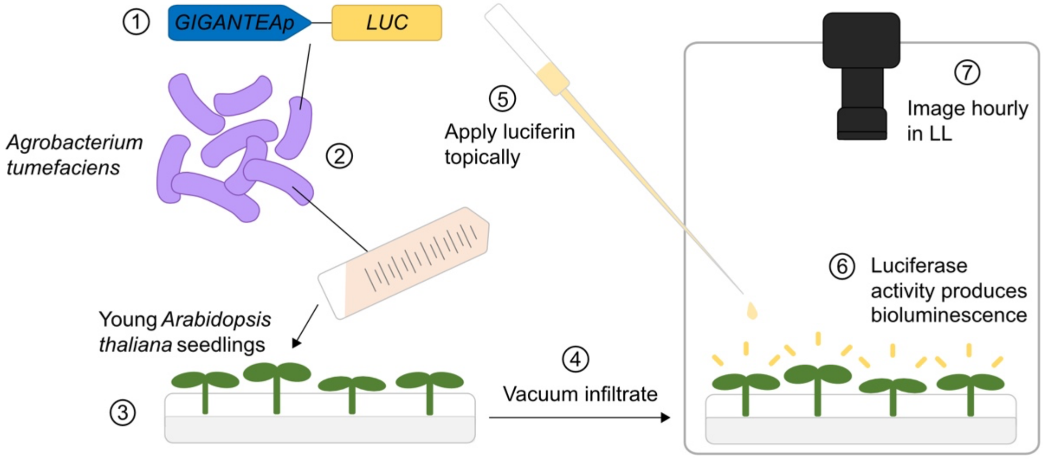


**Supplementary Figure S1: Overview of transient luciferase imaging for measuring Arabidopsis circadian rhythms.** The *GIGANTEAp:LUC2* (*GIp:LUC2*) construct is transiently transformed to Arabidopsis seedling clusters via *Agrobacterium tumefaciens*-mediated transformation. Luciferin is applied to transformed seedlings, and rhythms of clock-driven luminescence are imaged hourly in continuous light.

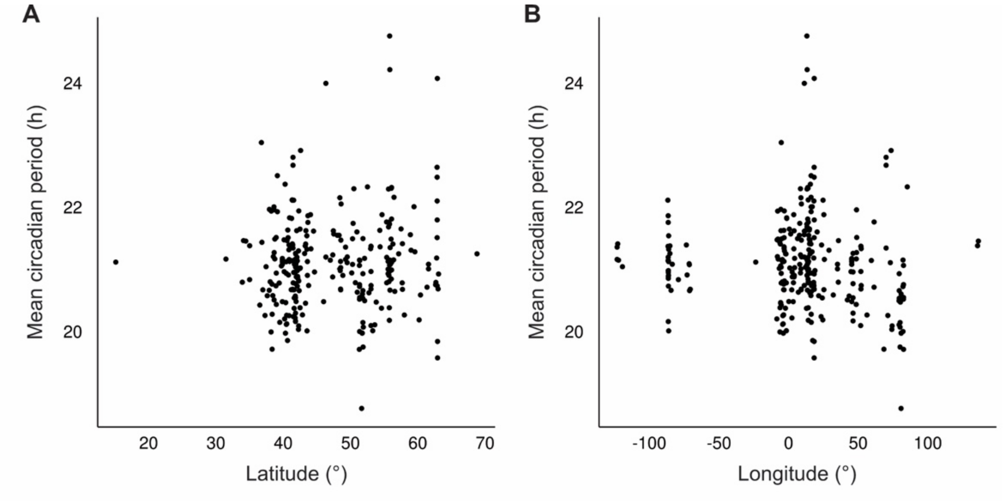


**Supplementary Figure S2: Relationship between mean circadian period of *Gip:LUC2* activity and geographical variables.** Each point represents an individual accession. **A,** Mean circadian period versus latitude of origin. **B,** Mean circadian period versus longitude of origin.

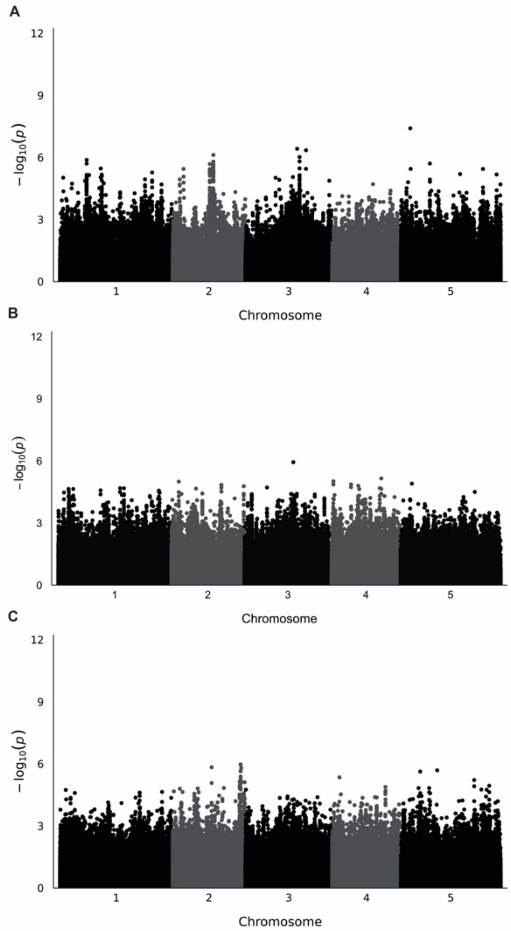


**Supplementary Figure S3: GWAS of circadian phase and RAE. A,** GWAS -$\log_{10}(P)$ values of association of SNPs with circadian phase. **B,** GWAS -$\log_{10}(P)$ values of association of SNPs with Z-score normalised amplitude. **C,** GWAS -$\log_{10}(P)$ values of association of SNPs with RAE. All -$\log_{10}(P)$ values fell below the FDR-adjusted significance threshold for all traits.

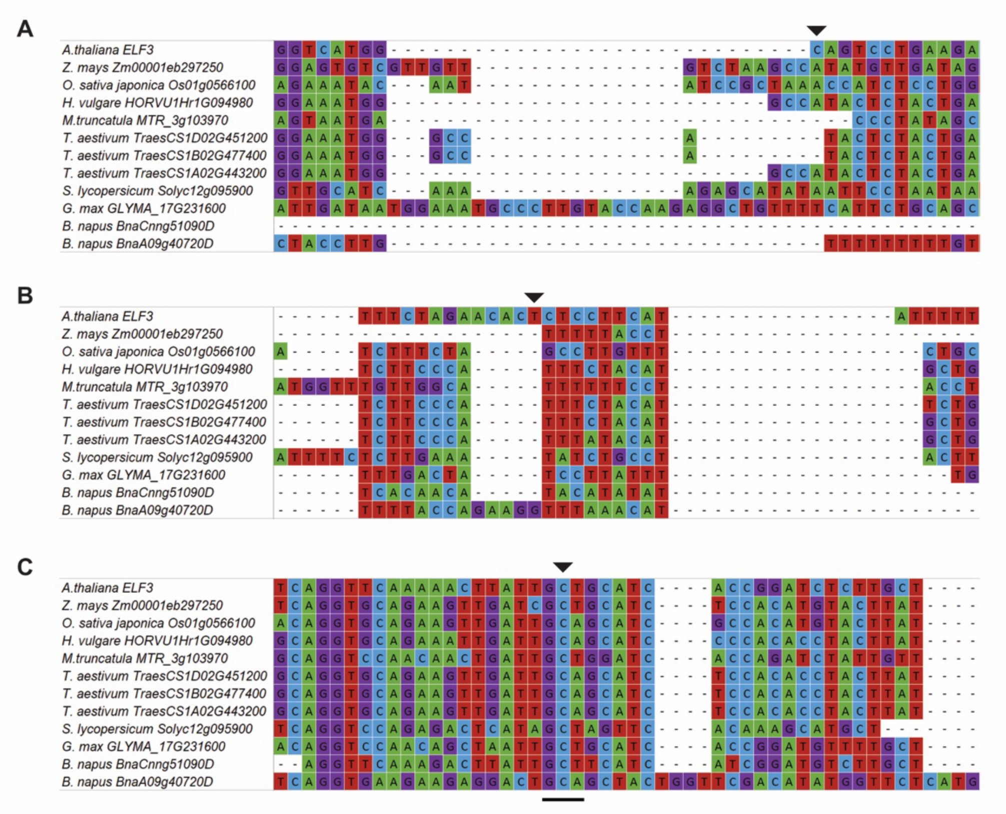


**Supplementary Figure S4: Conservation of *ELF3* SNP1-3 across plants.** *ELF3* genomic sequences were obtained from multiple plant species and aligned. The positions of SNP1 **(A)**, SNP2 **(B)** and SNP3 **(C)** are highlighted by black arrows. Black line shows position of GCT and GCA alanine codons which are conserved across plants. *ELF3* sequences were sourced from *Arabidopsis thaliana*, *Zea mays*, *Oryza sativa*, *Hordeum vulgare*, *Medicago truncatula*, *Triticum aestivum*, *Solanum lycopersicum*, *Glycine max* and *Brassica napus*.

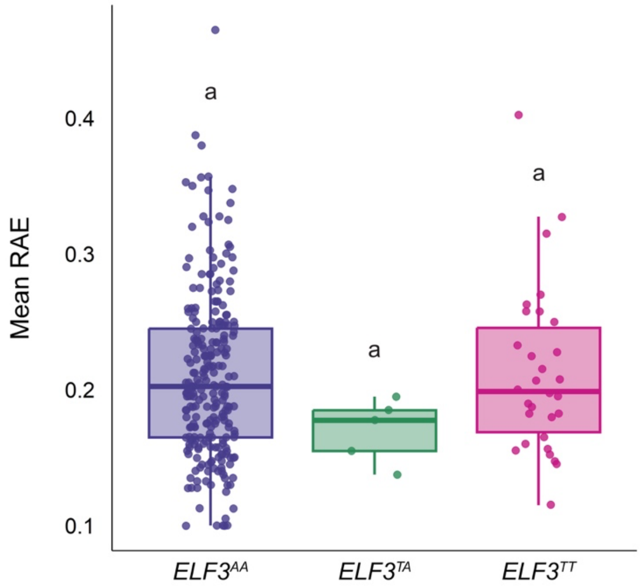


**Supplementary Figure S5: ELF3 polymorphisms are not associated with differences in robustness of rhythms.** Mean relative amplitude error (RAE) of rhythms of *Gip:LUC2* actiivity of Arabidopsis accessions grouped by *ELF3* haplogroup. Different letters indicate significant differences as determined by one-way ANOVA followed by Tukey's HSD; $P < 0.05$.

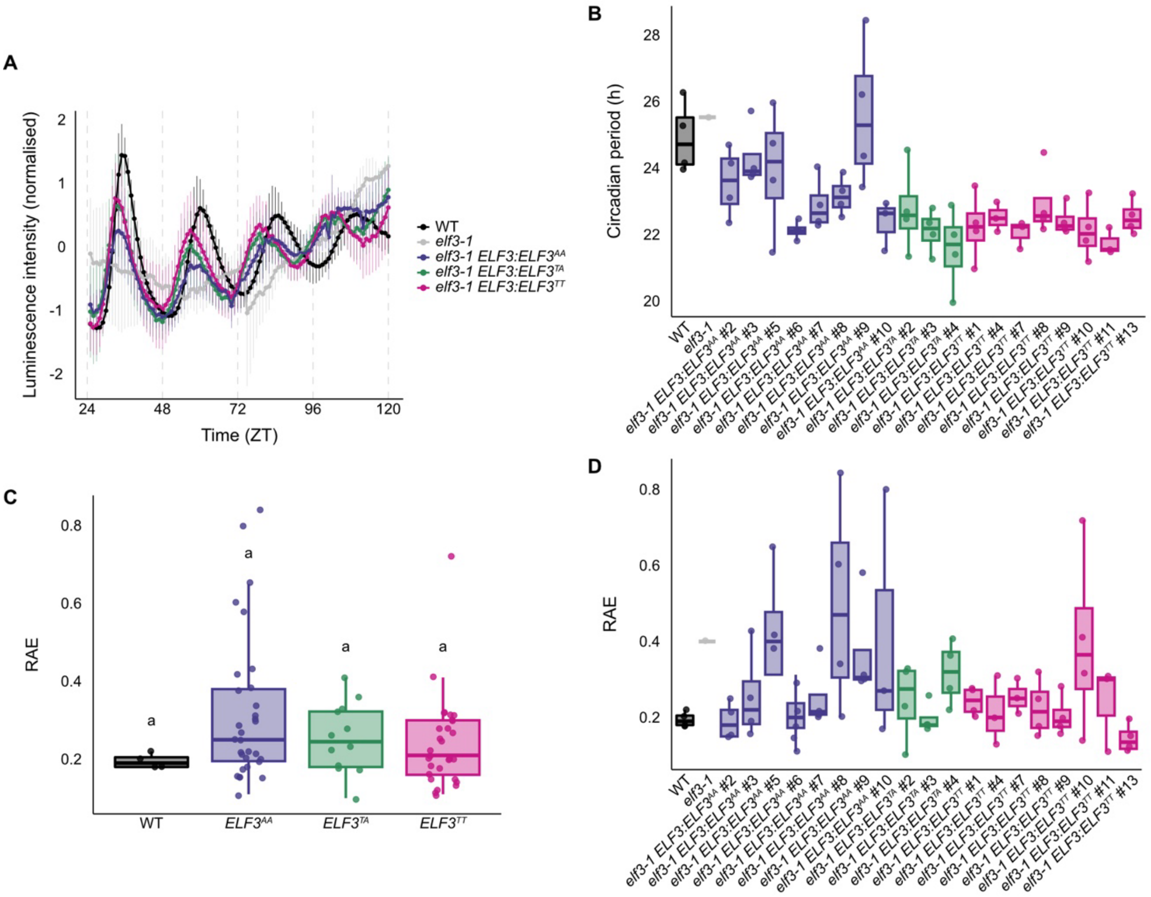


**Supplementary Figure S6: Circadian rhythms of transgenic *elf3-1* plants complemented with *ELF3* alleles from three haplotypes.** Circadian rhythms of *elf3-1* complementation lines and Col-0 (WT) and *elf3-1* were measured via seedling transformation of *GIp:LUC2* reporter constructs. **A,** Normalised luminescence intensity over a circadian time series. Haplotype groupings represent multiple transgenic lines ($ELF3^{AA}$: 8 lines, $ELF3^{TA}$: 3 lines, $ELF3^{TT}$: 8 lines). For individual lines, n=4. Error bars represent ±sd, **B,** Circadian period values of individual *elf3-1* complementation lines, n=4. **C,** RAE of *elf3-1* complementation lines. Haplotype groupings represent multiple transgenic lines ($ELF3^{AA}$: 8 lines, $ELF3^{TA}$: 3 lines, $ELF3^{TT}$: 8 lines). For individual lines, n=4. Different letters indicate significant differences as determined by one-way ANOVA followed by Tukey's HSD; $P < 0.05$. **D,** RAE values of individual *elf3-1* complementation lines, n=4.

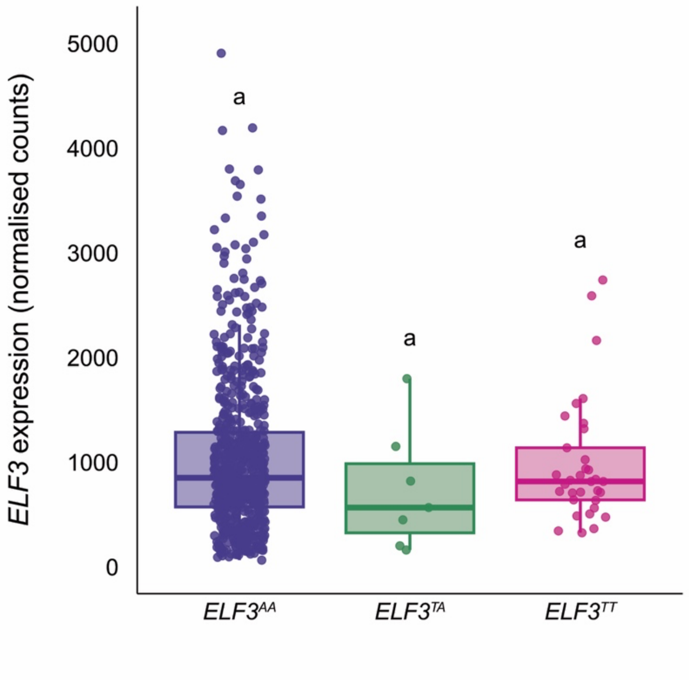


**Supplementary Figure S7: *ELF3* expression does not differ between haplogroups.** *ELF3* (*ELF3α* main isoform) transcript data for 728 accessions were sourced from Kawakatsu *et al.* (2016). Different letters indicate significant differences as determined by one-way ANOVA followed by Tukey's HSD; $P < 0.05$.

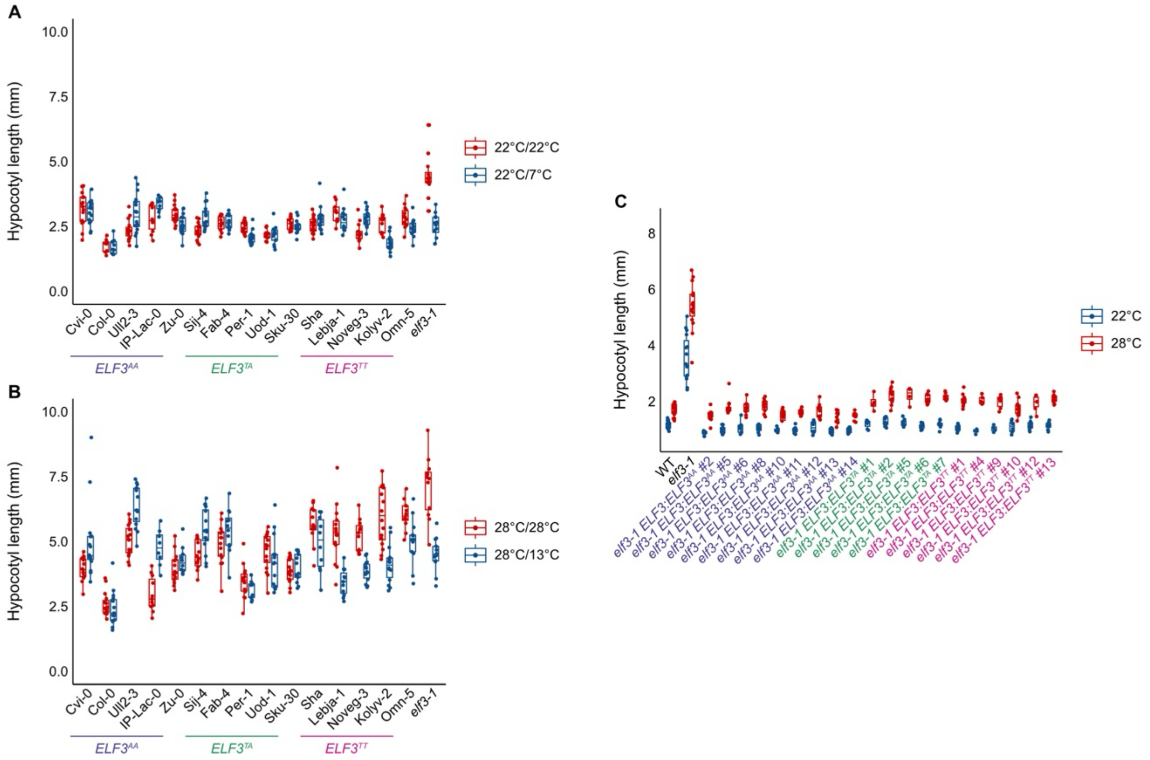


**Supplementary Figure S8: *ELF3* polymorphisms are associated with differences in hypocotyl elongation at elevated temperature.** Quantification of hypocotyl elongation in different temperature regimes. **A,B,** Hypocotyl elongation of naturally occurring Arabidopsis accessions representing each of the three *ELF3* haplogroups (*ELF3*$^{AA}$: Col-0, IP-Lac-0, Zu-0, Ull2-3, Cvi-0; *ELF3*$^{TA}$: Sku-30, Fäb-2, Uod-1, Per-1, Sij-4; *ELF3*$^{TT}$: Omn-5, Kolyv-2, Lebja-1, Noveg-3 and Shahdara). **A,** Constant control temperature (22°C: 22°C) or variable control temperature (22°C: 7°C). **B,** Constant elevated temperature (28°C: 28°C) or variable elevated temperature (28°C: 13°C). Haplogroup groupings in Figure 5A-C are comprised of the individual accessions shown here. **C,** Hypocotyl elongation of transgenic *elf3-1* plants complemented with *ELF3* alleles at constant temperatures. Haplotype groupings in Figure 5D are comprised of the individual lines shown here**.** For all experiments, n=16.

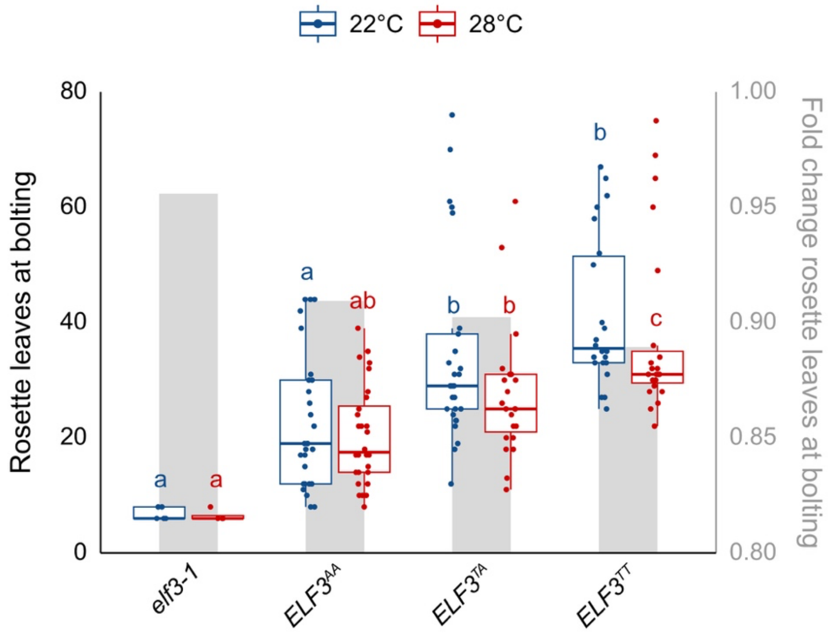


**Supplementary Figure S9: *ELF3* polymorphisms are associated with differences in flowering time at elevated temperature.** Number of rosette leaves at bolting (left axis) and fold change of mean number of rosette leaves at bolting (grey boxes, right axis) between constant 22°C and 28°C for *elf3-1* and natural accessions (*ELF3*$^{AA}$: Col-0, IP-Lac-0, Zu-0, Ull2-3, Cvi-0; *ELF3*$^{TA}$: Sku-30, Fäb-2, Uod-1, Per-1, Sij-4; *ELF3*$^{TT}$: Omn-5, Kolyv-2, Lebja-1, Noveg-3 and Shahdara). Letters indicate significant differences within each temperature condition as determined by one-way ANOVA followed by Tukey's HSD; $P < 0.05$. n=6 for each accession.
